# Identification of biological processes and signaling pathways for the knockout of REV-ERB in mouse brain

**DOI:** 10.1101/2021.11.22.469579

**Authors:** Jing Li, Wei Wang, Hanming Gu

**Affiliations:** SHU-UTS SILC School, Shanghai University, Shanghai, China

## Abstract

REV-ERB is an orphan nuclear receptor that is widely expressed in the brain and inhibits transcriptional activities. A variety of genes affect the activity and expression of REV-ERB. In this study, our objective is to identify significant signaling pathways and biological processes in the knockout of the REV-ERB mouse brain. The GSE152919 dataset was originally created by using the Illumina HiSeq 4000 (Mus musculus). The KEGG and GO analyses suggested that biological processes “PPAR signaling”, “Hippo signaling”, and “Hypertrophic cardiomyopathy (HCM)” are mostly affected in the knockout of REV-ERB. Furthermore, we identified a number of genes according to the PPI network including NPAS2, CRY2, BMAL1, and CRY1 which were involved in the lack of REV-ERB in the brain. Therefore, our study provides further insights into the study of circadian clocks.

## Introduction

The circadian clocks control the behavior and physiology of living organisms according to the external environment^1, 2^. The core circadian clocks regulate transcriptional and physiological rhythms which form a transcriptional-translational feedback loop^3^. The core circadian clocks contain transcriptional activators Bmal1/ClOCK which activates their repressor proteins such as PER, CRY, and REV-ERB^4^. The circadian clocks control various cellular processes such as metabolism, inflammation, and mitochondrial homeostasis^5–9^. Clocks’ function and cycles of energy metabolism are closely and reciprocally linked^10–12^. The disruption of clocks leads to metabolic diseases such as type 2 diabetes and heart diseases^13^.

REV-ERBα is a nuclear reporter protein that is directly mediated by BMAL1^14^. REV-ERBα is also a transcriptional repressor that restrains Bmal1 expression and potential downstream genes at specific sites within the genome^15^. Given that REV-ERBα locates in the nucleus, it becomes a potential drug target that can be regulated by small-molecule agonists and antagonists^16^. Recent reports showed that REV-ERBα is one of the key regulators in mediating the energy metabolism^17^. The REV-ERBα mice depicted remarkable changes in the homeostasis of carbohydrate and lipid, which displayed an up-regulation of lipid accumulation and storage^18^. REV-ERBα is regulated by BMAL1/CLOCK heterodimers through transactivation and posttranslational protein degradation^17^. REV-ERBα indicates circadian rhythmic activity, which competes and binds with the RORE sites (AGGTCA hexamer with a 5′ A/T-rich sequence) of ROR proteins^19^. REV-ERBα knockout mice exhibited significantly changed cortical resting-state functional connectivity, which was found in neurodegenerative models^20^.

In our study, we evaluated the effects of knockout of REV-ERB in the suprachiasmatic nucleus (SCN) during the nighttime by analyzing the RNA sequence data. We identified a number of DEGs and the potential affected biological processes. We also performed the gene functional enrichment and constructed the protein-protein interaction (PPI) network for finding the potential interacting proteins. These important genes and biological processes could provide efficient guidance on drug development.

## Methods

### Data resources

Gene dataset GSE152919 was collected from the GEO database. The data was created by using the Illumina HiSeq 4000 (Mus musculus) (Institute for Diabetes, Obesity, and Metabolism, University of Pennsylvania Perelman School of Medicine, Philadelphia, PA19104-5160, US). The analyzed dataset includes 5 WT and 4 REV-ERB KO at CT16.

### Data acquisition and preprocessing

The data were conducted by the R package. We used a classical t-test to identify DEGs with P<.05 and fold change ≥1.5 as being statistically significant^21, 22^.

The Kyoto Encyclopedia of Genes and Genomes (KEGG) and Gene Ontology (GO) analyses KEGG and GO analyses of DEGs in this study were conducted by the Database for Annotation, Visualization, and Integrated Discovery (DAVID) (http://david.ncifcrf.gov/). P<.05 and gene counts >10 were considered statistically significant.

### Protein-protein interaction (PPI) networks

The Molecular Complex Detection (MCODE) was used to construct the PPI networks. The significant modules were created from constructed PPI networks. The pathway enrichment analyses were performed by using Reactome (https://reactome.org/), and P<0.05 was used as the cutoff criterion.

## Results

### Identification of DEGs in WT and REV-ERB KO

Since REV-ERB showed higher expression during the night in comparison with daytime, we analyzed the DEGs from the WT and REV-ERB KO mice at Zeitgeber times (ZT) 16^23^. A total of 228 genes were identified to be differentially expressed with the threshold of P<0.05. The up- and down-regulated genes for WT and REV-ERB KO samples were depicted by the heatmap and volcano plot (Figure 1). Among them, the top ten DEGs were selected and listed in Table 1.

**Table 1.**

| **Entrez gene** | **Gene Symble** | **Fold-change** | **Regulation** |
| --- | --- | --- | --- |
| Top 10 down-regulated DEGs |  |  |  |
| 170722 | Nxf7 | -2.914409539 | Down |
| 84063 | Kirrel2 | -2.408625265 | Down |
| 255349 | Tmem211 | -2.079681186 | Down |
| 74879 | 4930461G14Rik | -1.953485978 | Down |
| 51458 | Rhcg | -1.909884329 | Down |
| 64833 | Acot10 | -1.623019882 | Down |
| 102466639 | Mir6385 | -1.383534957 | Down |
| 73040 | 2900052N01Rik | -1.135864344 | Down |
| 4240 | Mfge8 | -1.127675784 | Down |
| 70393 | 2210416O15Rik | -1.0711531 | Down |
| Top 10 up-regulated DEGs |  |  |  |
| 26 | Aoc1 | 5.441097014 | up |
| 84033 | Obscn | 5.192832086 | up |
| 6524 | Slc5a2 | 3.785831939 | up |
| 2173 | Fabp7 | 3.745998775 | up |
| 83483 | Plvap | 2.931300223 | up |
| 23831 | Car14 | 2.885290644 | up |
| 594855 | Cplx3 | 2.611357879 | up |
| 79748 | Lman1l | 2.581577124 | up |
| 83716 | Crispld2 | 2.508233633 | up |
| 329271 | C230024C17Rik | 2.398221442 | up |

**Figure 1.**
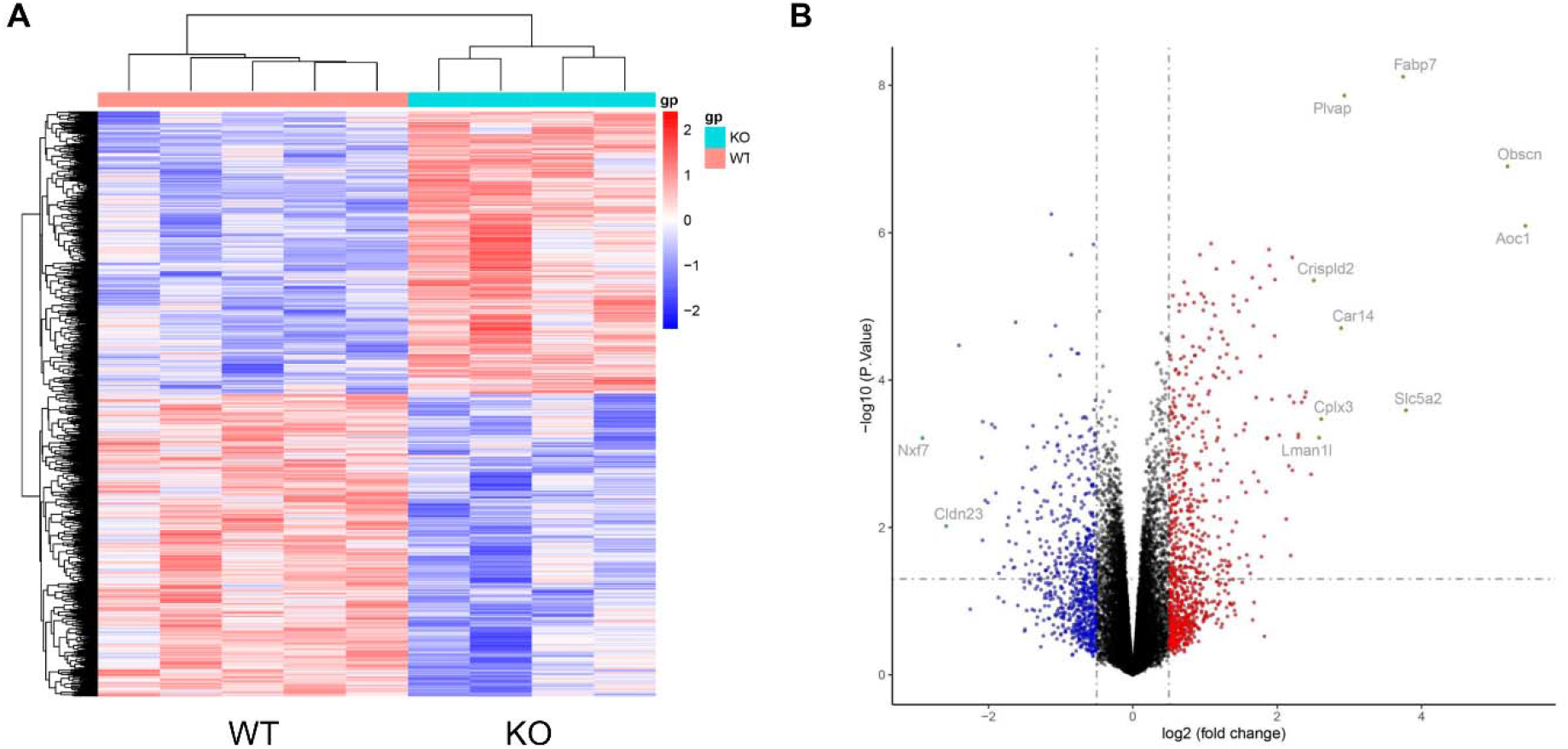
Heatmap and volcano plot between WT and REV-ERB KO. (A) Heatmap of significant DEGs between WT and REV-ERB KO. Regularized matrix was generated using the R package. Significant DEGs were used to create the heatmap. (B) Volcano plot for DEGs between WT and REV-ERB KO. The most significant genes are highlighted grey dots and gene symbols marked.

### Enrichment analysis of DEGs in WT and REV-ERB KO

To further understand the biological roles of REV-ERB, we performed KEGG and GO enrichment analysis (Figure 2). The top five significant KEGG pathways were analyzed including “ Circadian rhythm”, “Oxytocin signaling pathway”, “Hippo signaling pathway”, “Hypertrophic cardiomyopathy (HCM)”, and “PPAR signaling pathway”. We identified the top ten MF categories of GO including “Nucleoside−triphosphatase regulator activity”, “Enzyme activator activity”, “GTPase regulator activity”, “Metal ion transmembrane transporter activity”, “amide binding”, “GTPase activator activity”, “Calcium ion transmembrane transporter activity”, “Voltage−gated cation channel activity”, “calcium channel activity”, and “Voltage−gated calcium channel activity”.

**Figure 2.**
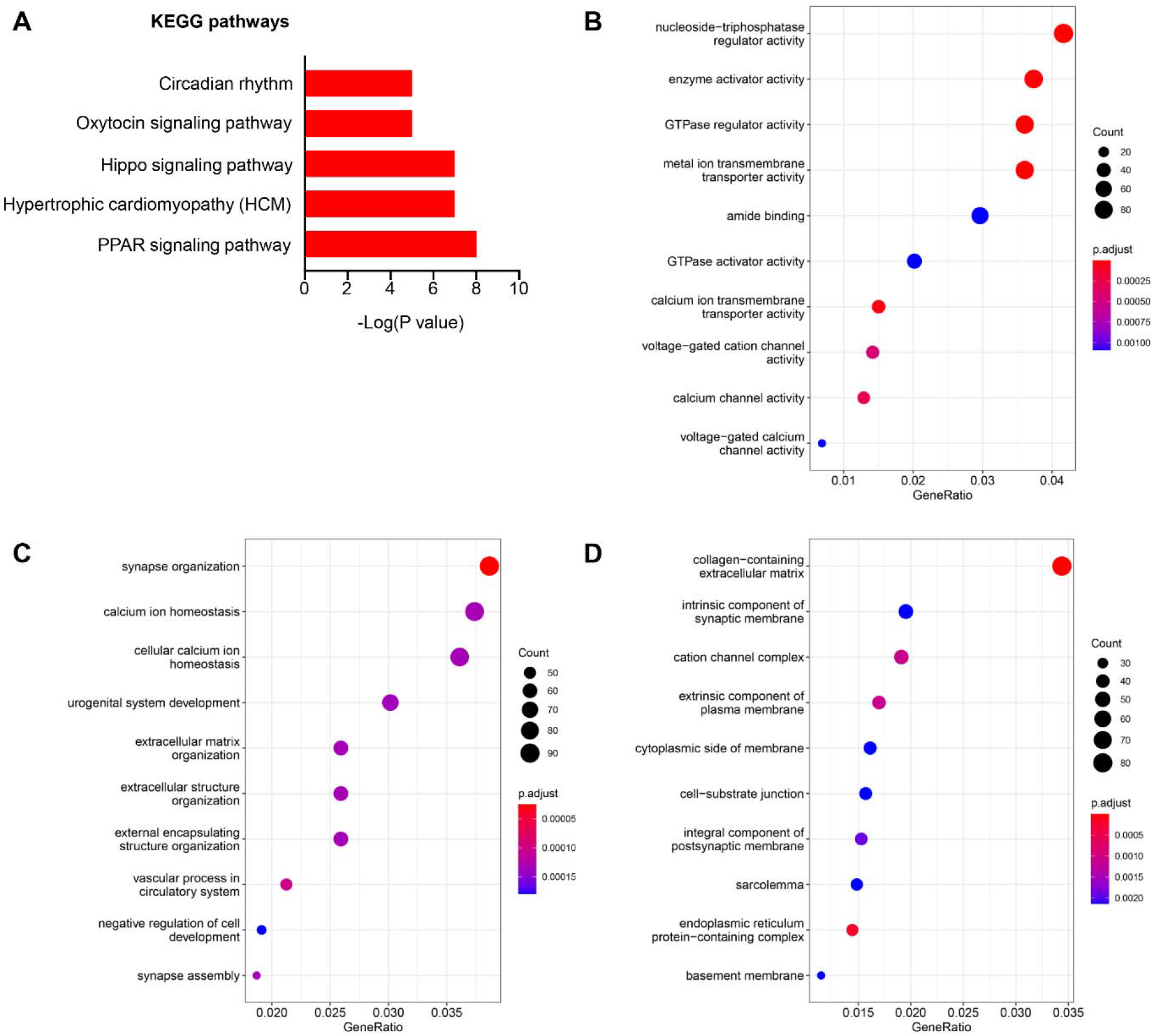
KEGG and GO analyses of DEGs between WT and REV-ERB KO. (A) KEGG analysis was performed by DAVID online tool. The significant terms were depicted. (B) Different colors represent biological processes (BP). (C) The molecular functions (MF) were analyzed by DAVID. (D) The cellular components (CC) were performed by DAVID.

Then, we identified the top ten BP categories of GO including “Synapse organization”, “Calcium ion homeostasis”, “cellular calcium ion homeostasis”, “Urogenital system development”, “Extracellular matrix organization”, “Extracellular structure organization”, “External encapsulating structure organization”, “vascular process in circulatory system”, “negative regulation of cell development”, and “synapse assembly”. We also identified the top ten CC of GO including “collagen−containing extracellular matrix”, “Intrinsic component of synaptic membrane”, “cation channel complex”, “extrinsic component of plasma membrane”, “cytoplasmic side of membrane”, “cell−substrate junction”, “integral component of postsynaptic membrane”, “sarcolemma”, “ndoplasmic reticulum protein−containing complex”, “basement membrane”.

### PPI network analysis in WT and REV-ERB KO

The PPI network was created to explore the relationships of DEGs affected by REV-ERB. The criterion of combined score >0.4 was set to construct the PPI by using the 127 nodes and 201 edges. The top ten genes with the highest degree scores are shown in Table 2. The top two significant modules were selected to depict the functional annotation (Figure 3). We also analyzed the DEGs and PPI with the Reactome analysis tools.

**Table 2.** Top ten genes demonstrated by connectivity degree in the PPI network.

| Gene Symbol | Gene title | Degree |
| --- | --- | --- |
| NPAS2 | Neuronal PAS domain protein 2 | 16 |
| CRY2 | Cryptochrome circadian regulator 2 | 16 |
| ARNTL | aryl hydrocarbon receptor nuclear translocator like | 14 |
| CRY1 | cryptochrome circadian regulator 1 | 14 |
| PER3 | period circadian regulator 3 | 13 |
| BHLHE41 | basic helix-loop-helix family member e41 | 13 |
| NR1D2 | nuclear receptor subfamily 1 group D member 2 | 13 |
| NR1D1 | nuclear receptor subfamily 1 group D member 1 | 13 |
| NFIL3 | nuclear factor, interleukin 3 regulated | 12 |
| TEF | TEF transcription factor | 12 |

**Figure 3.**
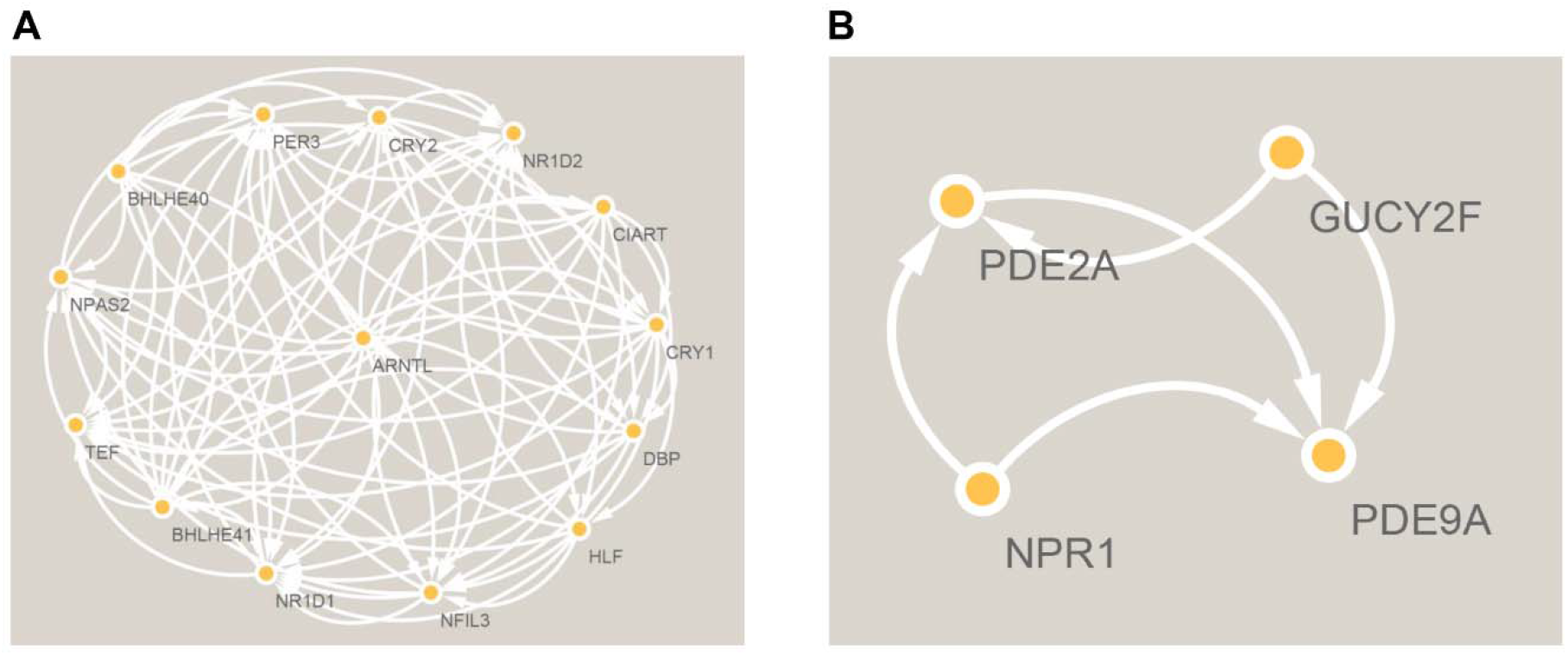
The PPI network analysis of DEGs between WT and REV-ERB KO. The 227 DEGs were input into the STRING database for PPI network analysis. Cluster 1 (A) and cluster 2 (B) were constructed by MCODE in Cytoscape.

The Reactome map showed the most biological functions affected by the knockout of REV-ERB (Figure 4). We also identified the top ten signaling pathways including “Circadian Clock”, “BMAL1:CLOCK, NPAS2 activates circadian gene expression”, “Transcriptional activation of cell cycle inhibitor p21”, “Transcriptional activation of p53 responsive genes “, “Heme signaling”, “TFAP2 (AP-2) family regulates transcription of cell cycle factors”, “TP53 Regulates Transcription of Genes Involved in G1 Cell Cycle Arrest”, “Sodium-coupled phosphate cotransporters”, “Extracellular matrix organization”, and “RUNX3 regulates CDKN1A transcription” (Supplemental Table S1).

**Figure 4.**
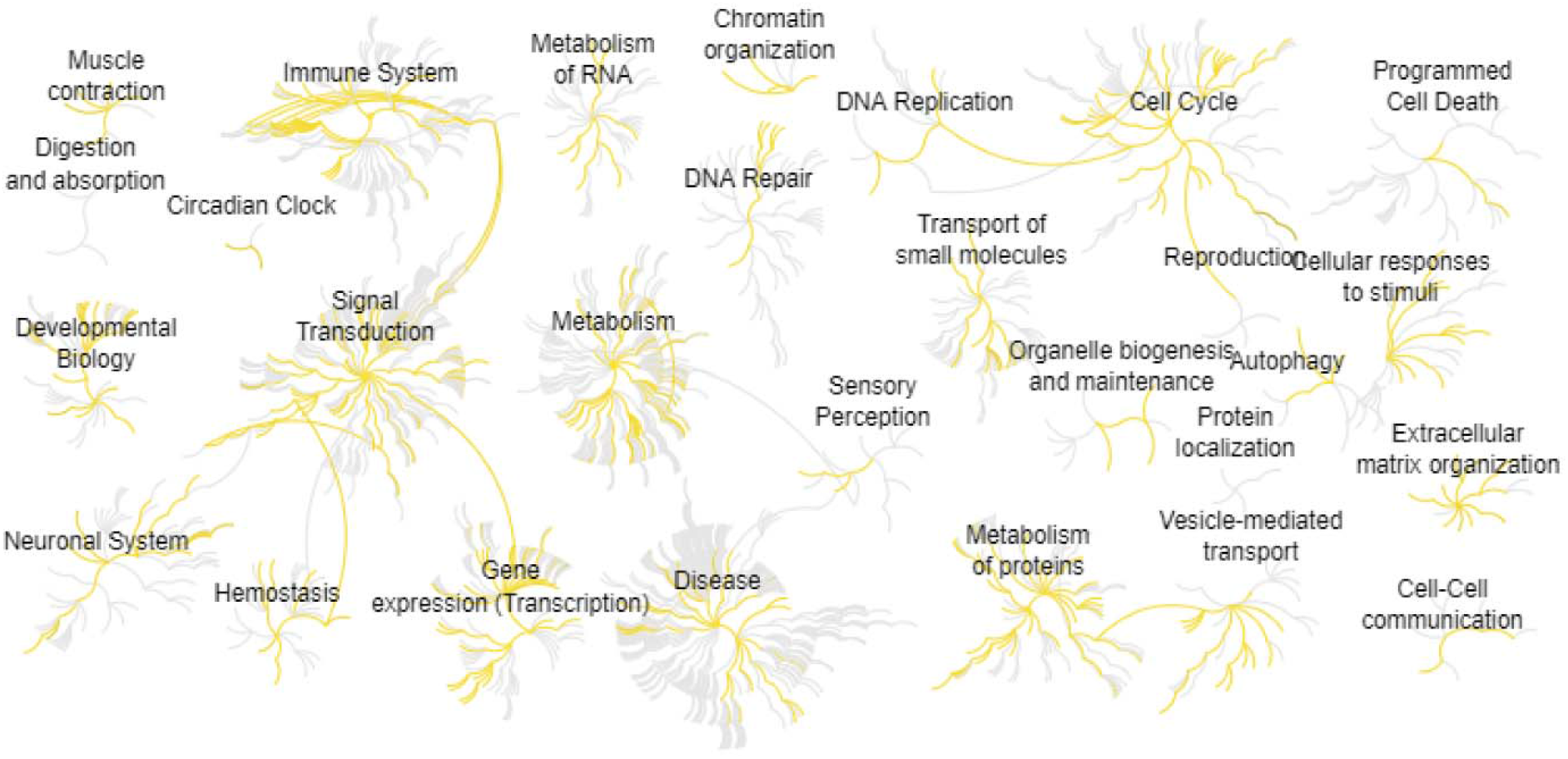
Reactome diagram representation of the significant biological processes of the protein elements identified between WT and REV-ERB KO.

## Discussion

As one of the transcriptional repressors, REV-ERB can inhibit gene transcription by recruiting co-factors nuclear receptor co⍰repressor 1 (NCOR1) and histone deacetylase 3 (HDAC3)^24^. Given that REV-ERBα regulates the clock and metabolic genes, it is proposed as a drug target for treating metabolic syndromes such as obesity and diabetes^25^. Recent studies showed various roles of REV-ERBα including inflammatory diseases, cancers, and heart diseases^25^. Moreover, REV-ERBα and its ligands have been considered valuable pharmacological molecules^16^.

REV-ERB is important for the development of metabolic diseases^16^. In our study, the KEGG analysis showed PPAR signaling, Hippo signaling, and Hypertrophic cardiomyopathy (HCM) are the most affected biological processes during the knockout of REV-ERB at night. In the study by Coralie Fontaine, REV-ERBα can activate the PPAR and further drive the adipocyte differentiation^26^. Protein modification is an important step in regulating molecular activity under physiological and pathological conditions. REV-ERBα facilitates cytosolic and nuclear protein O-GlcNAcylation that can change the activity of YAP in the hippo signaling pathway^27^. Moreover, Lilei et al found REV-ERBα inhibits heart failure by repressing the transcription^28^. These findings are supported by our study. In the study of GO (BP analysis), we found that the nucleoside-triphosphatase regulator activity was the most affected process during the deficiency of REV-ERB. The core circadian clocks such as BMAL1 and CLOCK were located in the nucleus and regulated the transcriptional activities of target genes including NF-κB and RANKL to further regulate the downstream signaling pathways^29–31^. REV-ERBα was also located in the nucleus and repressed the activity of BMAL1^32^. It is suggested that REV-ERBα may affect the transcriptional activities through BMAL1. Interestingly, we also found that the knockout of REV-ERB can affect GPCR signaling pathways. GPCR and RGS signaling pathways play key roles in mediating the physiological and pathological processes such as metabolism^33, 34^, inflammation^35–39^, and tumorigenesis^40, 41^. It was found that REV-ERB forms complexes with NR2E3 to further regulate the expression of Guanine nucleotide-binding protein 1 (Gnat1)^42^. We also found that REV-ERB KO affects the synapse organization and assembly. Supportively, Tianpeng et al found the disorder of REV-ERBα inhibits GABAergic function and drives epileptic seizures in preclinical models^43^. Moreover, REV-ERB regulates the complement expression and microglial synaptic phagocytosis^44^.

In our study, the PPI analysis identified a number of critical genes that may affect the biological processes in REV-ERB KO mice. NPAS2 is highly expressed in the brain and can control the anxiety-like behavior and GABAA receptors^45^. Cry2 is one of the core components of circadian clocks which has been linked to depression in patients^46^. BMAL1 is a basic helix loop helix transcription factor that binds with its partner CLOCK or NPAS2 to control the circadian oscillations^47^. The circadian controlled pathways include PER, CRY, NR1D1, and other genes that underlie circadian oscillation of ER stress, molecule activity, and oxidant defenses^48–51^. Cry1 is also highly expressed in the brain that associates with PER, which leads to the inhibition of CLOCK-BMAL1 to further control the clock-controlled genes^52^. As a core circadian gene, Per3 can regulate the embryonic development of the cerebral cortex^53^. Bhlhe41 is required for the competitive fitness of alveolar macrophages and the knockout of Bhlhe41 inhibits the proliferation of macrophages^54^. Hui et al found NR1D2 can promote the progression of liver cancer by regulating the epithelial transition^55^. As a survival factor, NFIL3 can inhibit the FOXO-regulated gene expression in cancer^56^. TEF (thyrotroph embryonic factor) is an important factor of the PAR bZip members, which is expressed in the brain and is relevant to intractable epilepsy^57, 58^.

In summary, our study provided the insight into the knockout of REV-ERB in mouse brains. The PPAR signaling, Hippo signaling, and Hypertrophic cardiomyopathy (HCM) are the significant biological processes during the deficiency of REV-ERB in the brain. Our future studies will explore the upstream and downstream of the important processes. Our study may facilitate the research on circadian rhythms.

## Author Contributions

Jing Li, Wei Wang: Methodology and Writing. Hanming Gu: Conceptualization, Methodology, Writing- Reviewing and Editing.

